# The dynamics of wild *Vitis* species in response to climate change facilitate the breeding of grapevine and its rootstocks with climate resilience

**DOI:** 10.1101/2024.09.28.615618

**Authors:** Mengyan Zhang, Xiaodong Xu, Tianhao Zhang, Zhenya Liu, Xingyi Wang, Xiaoya Shi, Wenjing Peng, Xu Wang, Zhuyifu Chen, Ruoyan Zhao, Wenrui Wang, Yi Zhang, Zhongxin Jin, Yongfeng Zhou, Zhiyao Ma

## Abstract

Climate change poses significant challenges to agricultural suitability and food security due to the limited adaptability of domesticated crops, however, our understanding of their climate adaptability is still limited. In this study, we compare the potential distributional dynamics in response to future climate change between grapevine and its wild relatives (*Vitis* L.) based on ecological niche models (ENMs). We reveal that the annual mean temperature is the primary factor influencing the potential distribution of cultivated grapes (*V. vinifera* L.). By 2080, under RCP 8.5, suitable areas for wine and table grapes from *V. vinifera* are predicted to reduced by 15.04×10^5^ km² and 13.26×10^5^ km², respectively. The results suggest that grape cultivation, particularly for table grapes, will face serious threats from future climate change. In contrast, seventy percent of wild grapes are well able to withstand future climate conditions. For example, under RCP 8.5, North American wild grapes like *V. rotundifolia* and *V. labrusca*, as well as East Asian wild grapes such as *V. heyneana* and *V. davidii*, are projected to demonstrate significant adaptability. These wild grapes have the potential to provide rootstocks that could enhance the adaptability of cultivated grapes and serve as invaluable resources for breeding programs aimed at developing new cultivars better suited to withstand the environmental pressures of climate change. Our results not only predict potential suitable distribution areas of wild grapes in the future, but also emphasize the essential impact of these wild genetic resources on grape breeding for promoting adaptation to future climate change.

## Introduction

Global climate change has significant implications for food security and agricultural sustainability, affecting some crucial agricultural traits such as biodiversity, growth, and overall productivity of crops (David et al., 2011; Yadav 2015; Brett et al., 2016; Pironon et al., 2019; You et al., 2024; Yi Yanget al., 2024). Climate change may lead to shrink in the acreage of important crops as they may struggle to thrive in their traditional growing regions, with increasing temperature and rising sea levels (Muller et al., 2011; Tim Wheeler et al., 2013). For example, wheat is considered the most vulnerable crop in eastern Africa, with projections suggesting up to 72% reductions in current yields by the end of this century (Adhikari et al., 2015). This trend is not limited to wheat, the productivity of grain legumes will be affected by the shrinking and degradation of arable land due to global climate change (Vadez et al., 2012). Additionally, climate change is predicted to adversely affect major crops, especially in tropical regions such as Brazil, with maize production facing significant declines due to delayed rainy seasons and rising temperatures, highlighting an urgent need for adaptive agricultural strategies to ensure food security (Dhaliwal et al., 2022; Bigolin et al., 2024). These challenges underscore the importance of resilient agricultural practices to mitigate the effects of climate change on our food supply.

The crop wild relatives (CWRs), as compared with domesticated cultivars, has been shown to be reliance to global climate change, and could adapt to some harsh environment which is unsuitable for domesticated cultivars. In such a scenario, increasing agricultural plant diversity (agrobiodiversity) and the utilization of crop wild relatives becomes crucial in improving crop response to future climate challenges (Waha et al., 2018). Compared with domesticated cultivars, CWRs possess untapped genetic diversity and provide valuable traits that can be effectively utilized for crop improvement with genome editing techniques, such as disease resistance and drought tolerance, in the face of changing climatic environments (Zhang et al., 2017; Li et al., 2018; Fernie & Yan, 2019; Renzi et al., 2022). Increasing attention has focused on the role of CWRs on promoting climate adaptation of crops recently. It has been suggested that crops in sub-Saharan Africa, such as potato, squash, and finger millet, could be sustained by utilizing CWRs to response to future climate change (Pironon et al., 2019; Satori et al., 2022). Similarly, the global distribution and conservation status of Rice Wild Relatives (RWRs) have been evaluated, with strategies to ensure the resilience of rice genetic resources in the face of climate change (Lin et al., 2024). To overcome challenges, researchers are exploring a range of strategies. This includes the genomic tools are being developed to introduce these traits into common legumes, which are critical for global food security (Porch et al., 2013). Concurrently, the investigation of alternative varieties, rootstocks, or geographic relocations may be explored (Rogiers et al., 2022). In summary, the impact of global climate change on crops highlights the importance of using CWRs in agricultural strategies to enhance crop resilience.

Grapevine (*Vitis vinifera* L.) is one of the most important fresh fruits, and widely cultivated around the world for wine (Zhou et al., 2017; Yang et al., 2023; Xiao et al. 2023). However, viticulture is very sensitive to climate change, which significantly limits its suitable growing regions (Hannah et al., 2013; Morales-Castilla et al., 2020). Climate change affects grapevine growth by influencing phenological stages such as bud burst, flowering, and ripening (Yang Dong et al., 2023). Changes in temperature and precipitation can alter the timing of these stages, potentially impacting grapevine distribution and productivity (Van Leeuwen et al., 2019; Rogiers et al., 2022). Furthermore, climate change can also impact grape quality by influencing sugar accumulation, acidity levels, and phenolic compounds (De Orduna, 2010; Van Leeuwen & Destrac-Irvine, 2017). Higher temperatures during the growing season, as projected under future climate scenarios, may increase the accumulation of sugars, affecting the flavor profile and overall quality of the grapes (Mosedale et al., 2016). Additionally, climate change might have a detrimental impact on grapevines by increasing pest and disease pressure and decreasing crop yield (Caffarra et al., 2012). Wild grape relatives in grape genus (*V.* L.) are important genetic resources for developing rootstocks and grape breeding with desirable traits, including resistance to biotic and abiotic stresses, such as Pierce’s Disease, cold, drought, and salinity (Morales-Cruz et al., 2021; Wang et al., 2021; Wang et al., 2023; Cochetel et al., 2023; Zhang et al., 2024). Although the influence of environmental niches on the distribution and growth of wild grapes has attracted researchers’ attention (Rahimi et al., 2021; Aguirre□Liguori et al., 2022; Petitpierre et al., 2023; Ma et al., 2023; Morales-Cruz et al., 2023), the assessment of the potential of wild grape relatives to respond to future climate is still lacking.

In this study, we aim to (1) simulate the niche distribution area and investigate climate factors preferment of cultivated grape and its wild relatives; (2) evaluate the response of cultivated grape and its wild relatives to future climate change; (3) identify wild species with potential for breeding and as rootstocks to response to future climate change.

## Results

### Cultivated grapes (*V. vinifera* L. ssp. *vinifera*) distributional potential in response to future climate change

The environmental suitability of cultivated grapes (*V. vinifera* L.) was estimated to have decreased, based on existing distribution data using the maxent model. Understanding the response relationship between species distribution and environmental factors is crucial for evaluating the impact of climate change on plant populations, taking cultivated grapes (*V. vinifera* L.) as an example, under current climate conditions, the suitable distribution areas for *V. vinifera* are widely distributed across East Asian, Europe, North America and southern Australia, with green representing table grapes and blue representing wine grapes for clear distinction (Fig. 1a-f). However, under the RCP 4.5 and RCP 8.5 scenarios (Representative Concentration Pathways (RCPs) describe potential future trajectories of greenhouse gas concentrations, with RCP 4.5 and RCP 8.5 being two of the scenarios considered), the suitable distribution for *V. vinifera* is expected to be significantly reduced. This indicates the substantial potential impact that high-concentration greenhouse gas emissions could have on the geographical distribution of *V. vinifera*. Climate variables, such as mean annual temperature and extreme temperatures, have a significant negative impact on the spatial distribution of grapes. In this study, the Annual Mean Temperature (bio1) was identified as the primary factor influencing the potential distribution of *V. vinifera* (Fig. S1), which is followed by the Mean Temperature of the Coldest Quarter (bio11) and the Minimum Temperature of the Coldest Month (bio6). The Annual Mean Temperature (bio1) has been identified as a primary factor influencing the potential distribution of grapes. In order to clarify the suitability of grape growth under different temperatures, we analyze the impact of bio1 on both wild and cultivated grapes. The analysis indicates that these grape varieties show different temperature adaptability, with bio1 having a greater impact on cultivated grapes (Fig. S1). Wild grapes are usually more adaptable and can grow in various climates, including those with extreme temperatures (Fig. 2). These results emphasize the critical role of temperature in determining the distribution of *V. vinifera* and also point to the challenges it may encounter due to climate change.

**Fig. 1.**
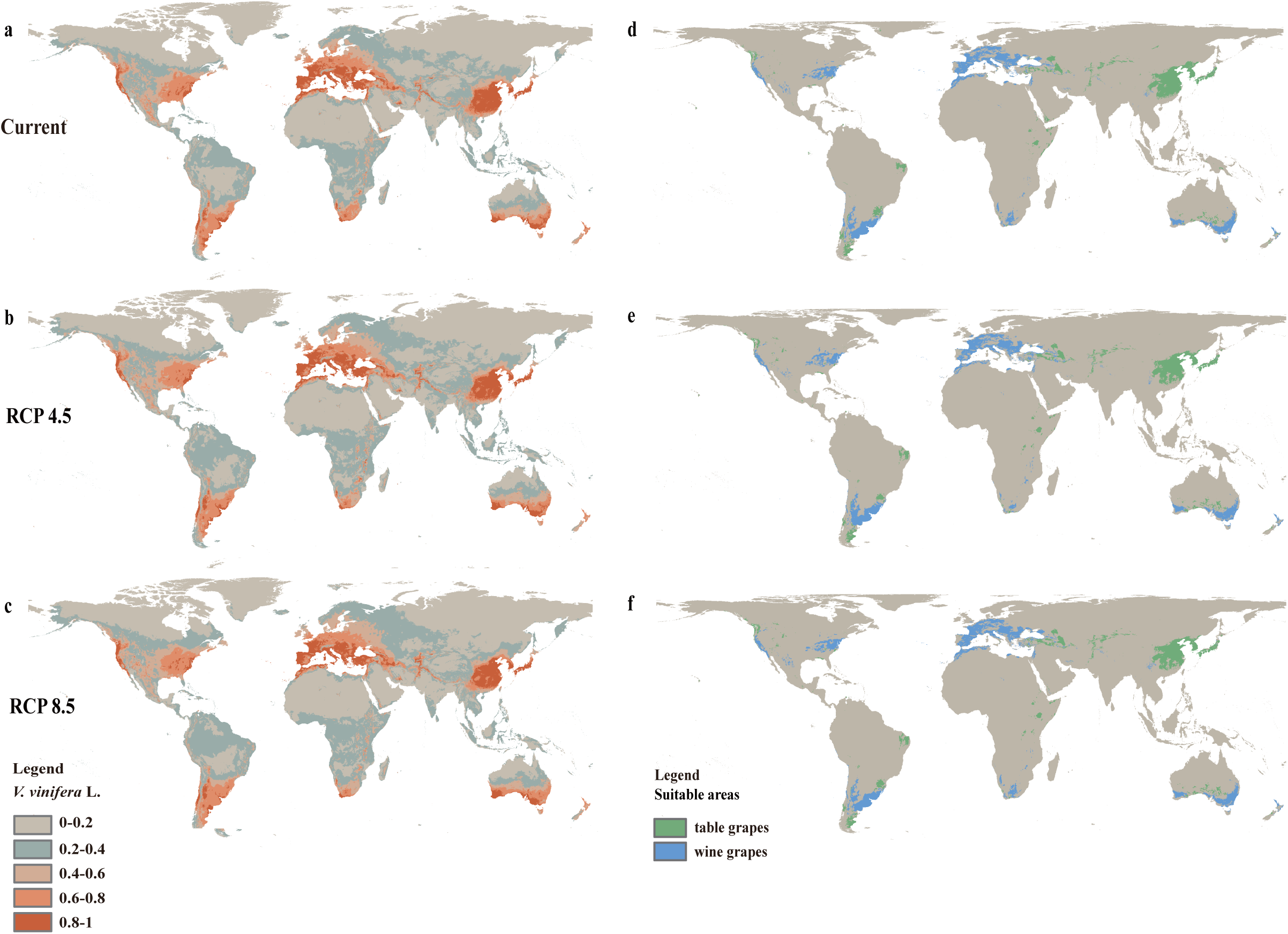
Suitable distribution ranges change of cultivated grapes (*V. vinifera* L.) under different climate change scenarios (a, d) current (1961–2000), (b, e) 2090 (2081–2100) RCP 4.5, (c, f) 2090 (2081–2100) RCP 8.5, green for table grapes and blue for wine grapes.

**Fig. 2.**
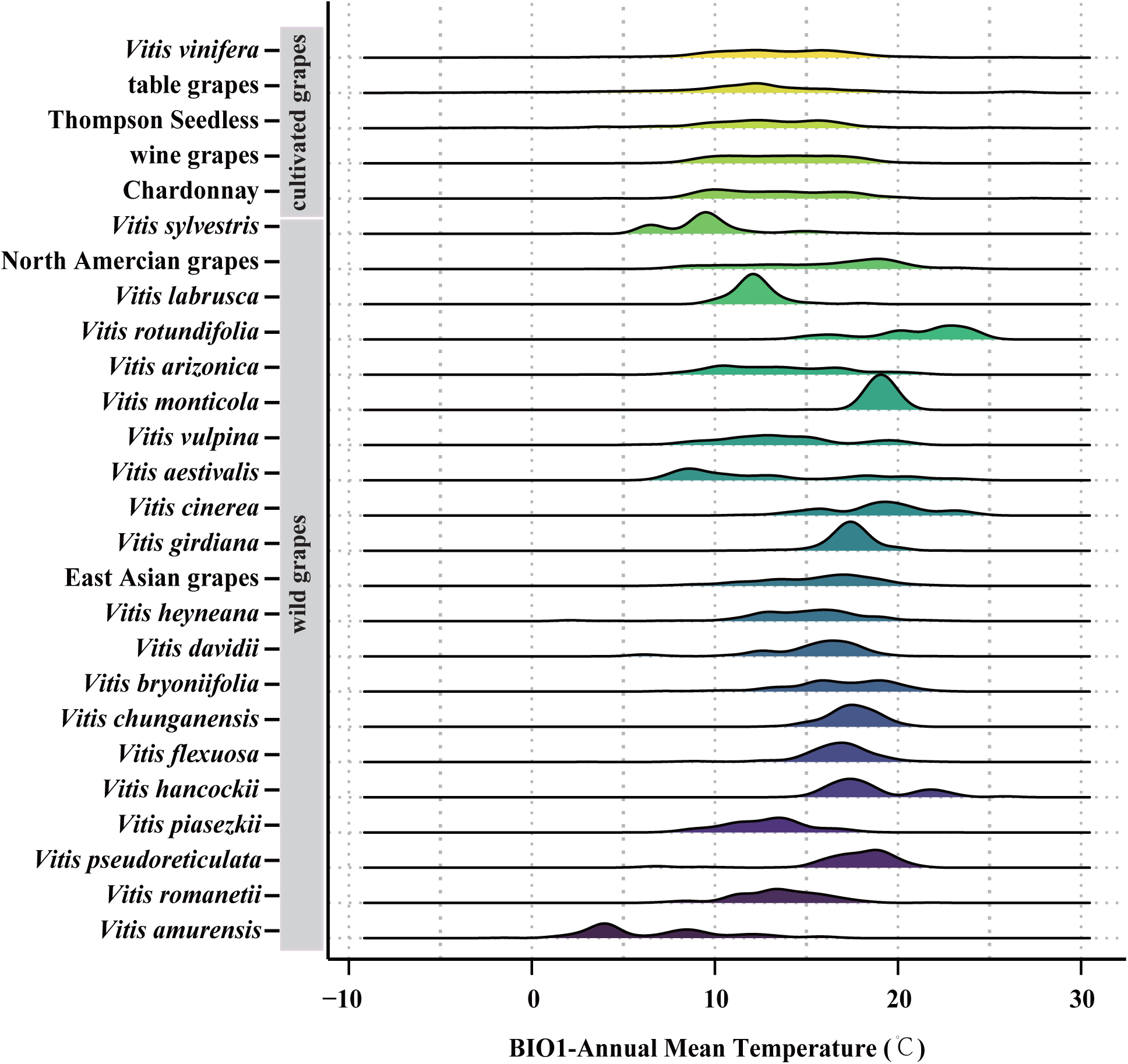
Temperature tolerance of cultivated grapes and wild grapes, the range of adaptability of different species to Annual Mean Temperature (bio1) the higher the peak, the better the growth condition of the variety may be within the temperature range.

Under RCP 8.5 scenario, projections indicate that the current suitable regions for *V. vinifera* will persist, encompassing an area of around 125.4×10^5^ km² (Table S1). The suitable areas for wine grapes and table grapes are expected to increase by 13.36×10^5^ and 19.12×10^5^ km² (Table S1). Despite overall expansion in the areas suitable for viticulture, the potential suitable areas for wine grapes decreased by 15.04×10^5^ km² and for table grapes, it decreased to 13.26×10^5^ km² (Table S1). The greater reduction for wine grapes compared to table grapes indicates that wine grapes may be more sensitive to climate change, or their suitable distribution conditions could be more negatively impacted.

In detail, current suitability is projected to be retained (Fig. 3c-d, refer to yellow color) in larger areas of current table grapes-producing regions, especially in China, although the model indicates declining suitability in the southern regions. Meanwhile, the suitability of Chile, the main table grapes producer in South America, is also declining (Fig. 3c-d, refer to red color), whereas the regions of northern China and northern Turkey are projected to become suitable in the future (Fig. 3c-d, refer to blue color). Specifically, current suitability is projected to be retained (Fig. 3e-3f, Fig. S2c-d, refer to yellow color) in many traditional wine-producing regions (such as France and Italy), under RCP 4.5 and RCP 8.5. Suitability is projected to decline (Fig. 3e-f; Fig. S2c-d, refer to red color) in Spanish regions and northeastern Australia (Queensland). Conversely, northern North American and two important non-Mediterranean wine-producing regions (non-Mediterranean Australia and New Zealand) are projected to expand in the future (Fig. 3e-f; Fig. S1-d, refer to blue color). As for Thompson Seedless, the most widely distributed table grapes, there is an observed trend of contraction in its suitable distribution range in southern China and Europe, as indicated by the red regions in Fig. S2a and b, conversely, in North America, the suitable areas for Thompson Seedless are expanding, as shown by the blue regions in the same figures. Chardonnay, the most widely distributed wine grapes, follows the same trend as wine grapes in terms of climate preferences. More interestingly, we explore that the suitability retained areas for wild grapes are comparable to 63% of those for cultivated grapes in the RCP 4.5 and 65% in the RCP 8.5. This suggests that some areas currently suitable for grape range may become unsuitable in the future.

**Fig. 3.**
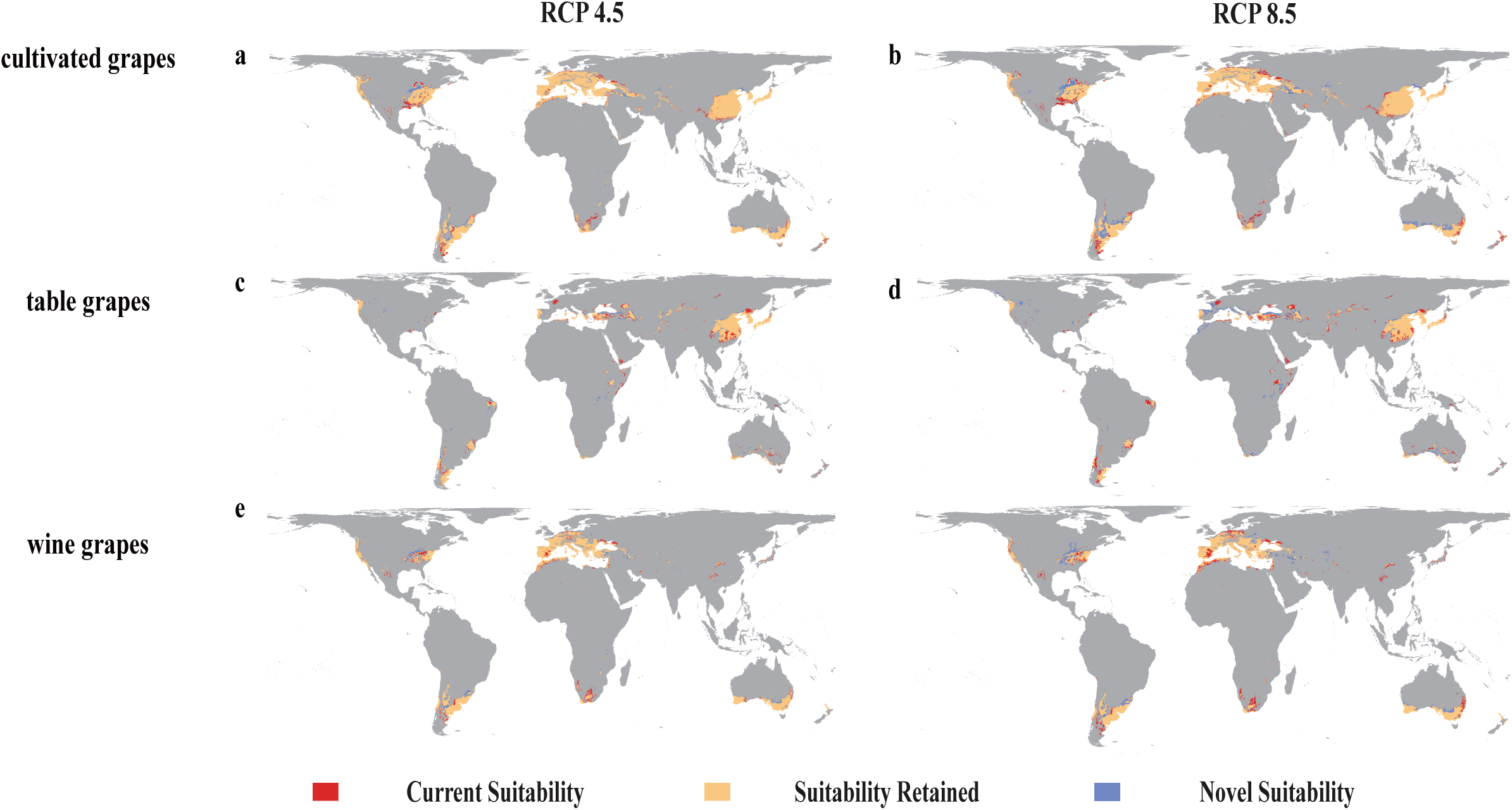
Maxent model projected bioclimatic suitability for cultivated grapes (*V. vinifera*), table grapes and wine grapes (a-b) cultivated grapes (*V. vinifera*), (c-d) table grapes, (e-f) wine grapes, with future projections for the years 2081–2100 under the RCP 4.5 and RCP 8.5 climatic scenarios. The projected bioclimatic suitability for Thompson Seedless and Chardonnay under the RCP 4.5 and RCP 8.5 scenarios can be found in the Supplementary Material (Fig. S2).

### Grape wild relatives in response to climate change

We compared European (*V. sylvestris*), North American, and East Asian wild grapes’ responses to climate change to identify species that may improve adaptability and resistance, the results show that current suitability is expected to be retained (Fig. 4a-f, refer to yellow color) in larger areas of current three wild grape regions. Among them, North American wild grapes have the largest area of 30.37×10^5^ km², followed by East Asian wild grapes and *V. sylvestris*, with 27.05×10^5^ km² and 23.88×10^5^ km² respectively (Table S1). As for *V. sylvestris*, declining suitability in the southeastern Europe, with a total decrease reaching 5.13×10^5^ km² (Table S1). In terms of the Expansion-Contraction Ratio, under the RCP 4.5 scenario, European wild grapes (*V. sylvestris*) exhibited a significant expansion trend in the future, increasing by 3.6%, with East Asian wild grapes following at 0.53%, while the suitability for North American wild grapes decreased by 0.32% (Fig. S3). As the climate gradually worsens, under RCP 8.5, the *V. sylvestris* Expansion-Contraction Ratio decreases to 1.8%. On the contrary, the East Asian wild grapes and North American wild grapes increase significantly, to 4.19% and 3.38% respectively (Fig. S3). This suggests that the East Asian wild grapes and North American wild grapes demonstrate a greater capacity to adapt to future adverse weather conditions compared to *V. sylvestris*.

**Fig. 4.**
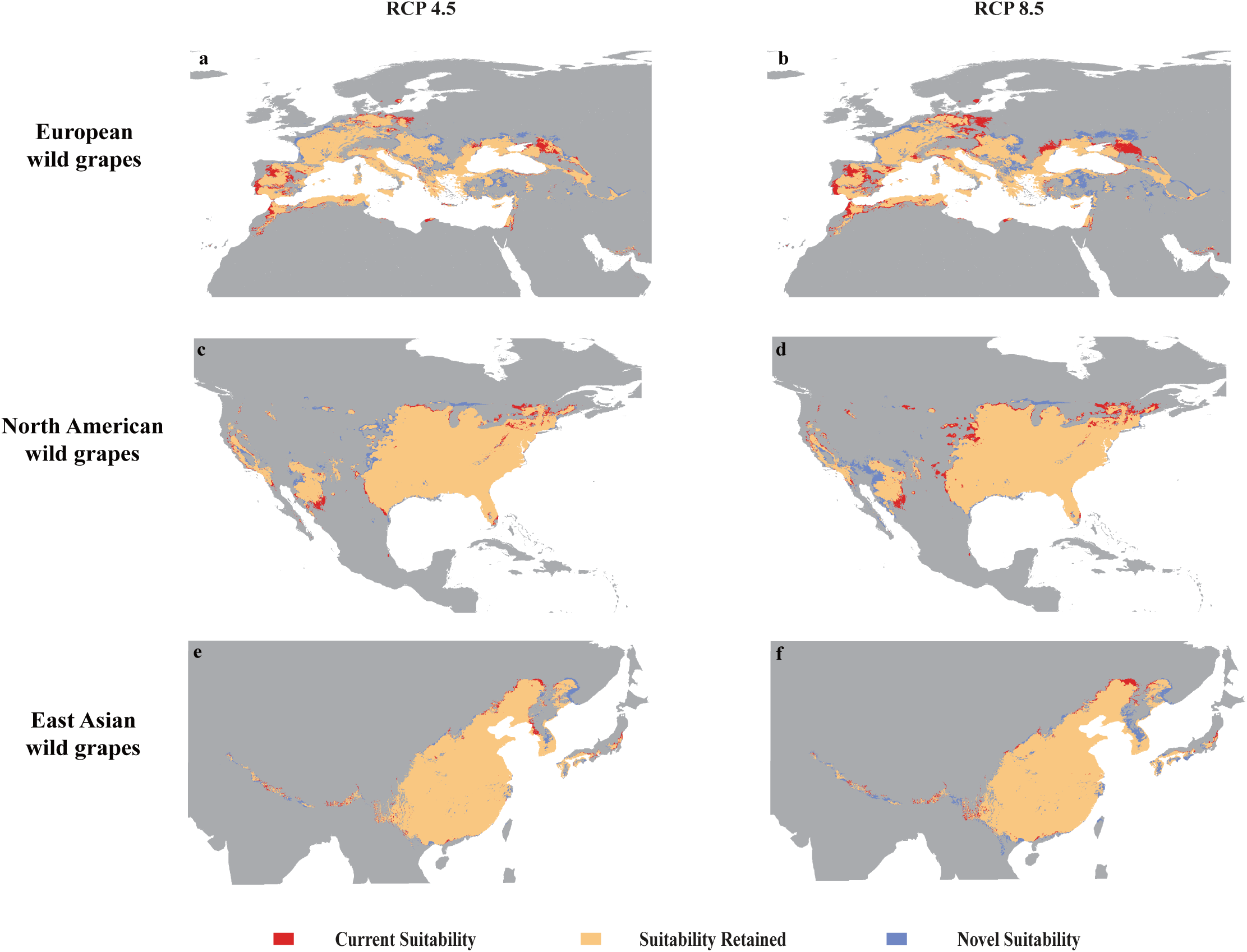
Maxent model projected bioclimatic suitability for European wild grapes (*V. sylvestris*), North American wild grapes, and East Asian wild grapes (a-b) European wild grapes (*V. sylvestris*), (c-d) North American wild grapes, (e-f) East Asian wild grapes, with future projections for the years 2081–2100 under the RCP 4.5 and RCP 8.5 climatic scenarios.

### Future suitability of North American wild grapes

An analysis of ten North American wild grapes distribution found that *V. rotundifolia* is expected to have the largest expansion of suitable areas in the future (Fig. 5b, refer to blue color), especially in the southeastern regions of North American. To understand these geographic shifts in more detail, we examine Expansion-Contraction Ratio analysis. Ensemble mean increases in suitable areas are 227% in North American in RCP 4.5, and 205% under RCP 8.5 (Fig. 5k). Followed by *V. labrusca* (Fig. 5a), large newly suitable areas are projected in regions of North American, accounting for 131% in RCP 4.5 and 118% under RCP 8.5 (Fig. 5k). They may be able to adapt to the warmer climate and changing patterns of precipitation. At the same time, the cultivation areas of *V. vulpina*, *V. aestivalis*, *V. monticola*, and *V. arizonica* have also shown an expanding trend, with respective increases of 11.49%, 9.28%, 7.88%, and 2.01%, while the distribution areas of *V. rupestris*, *V. riparia*, and *V. cinerea* are facing the risk of contraction. Under the RCP 8.5 scenario, the mean suitability decline rates were 12.91%, 1.91%, and 1.37% respectively, mainly in the middle and lower parts of North American (Fig. 5f, g, i, red). Unfortunately, with worsening climate conditions, the environmental requirements for *V. girdiana* can no longer be met in the North America, indicating that *V. girdiana* has become unsuitable for distribution there (Fig. 5j).

**Fig. 5.**
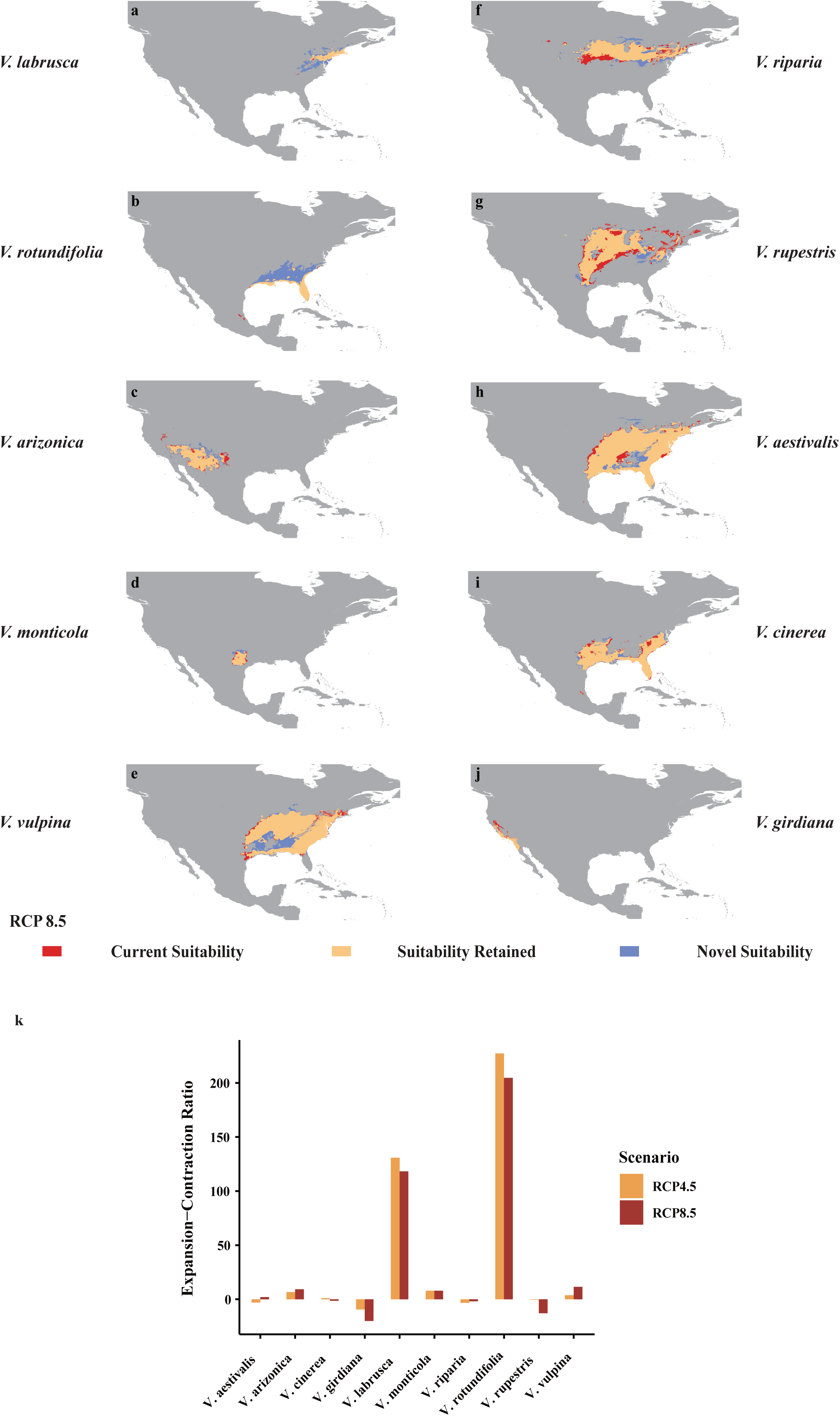
Maxent model projected bioclimatic suitability for North American wild grapes (a) *V. labrusca*, (b) *V. rotundifolia*, (c) *V. arizonica*, (d) *V. monticola*, (e) *V. vulpina*, (f) *V. riparia*, (g) *V. rupestris*, (h) *V. aestivalis*, (i) *V. cinerea*, (j) *V. girdiana*, with future projections for the years 2081–2100 under the RCP 8.5 climatic scenarios. (k) Net suitability changes for the distribution of North American wild grapes. Bar plots show Expansion-Contraction Ratio of change in area suitable for grape-growing regions projected by maxent model for RCP 8.5 (red) and RCP 4.5 (yellow).

### Future suitability of East Asian wild grapes

In terms of East Asian wild grapes, we analyzed the sensitivity and adaptability of ten grape species to climate change. Among them, *V. heyneana* shows an ensemble mean increase in the suitable areas of 49% in China under RCP 4.5, and 46% in RCP 8.5 (Fig. 6k). Large increases in ecological footprint are projected in eastern, central and western China (Fig. 6a, refer to blue color). *V. davidii* witnessed an opposite trend, compared with RCP 4.5, under RCP 8.5, the suitable areas increased by 2.3%, reaching 22%. *V. bryoniifolia*, *V. chunganensis*, *V. flexuosa*, *V. hancockii*, *V. piasezkii*, and *V. pseudoreticulata* are also showing a positive trend of expansion (Fig. 6k), the area is projected to experience an expansion ranging from 0.89% to 8.11% under the RCP 4.5, and between 3.89% to 8.65% under the RCP 8.5, attributed to their adaptability to environmental changes. In contrast, the suitable areas for *V. amurensis* are anticipated to contract. Under the RCP 4.5 scenario, the suitable areas are projected to decrease by 7.91%, while under the RCP 8.5 scenario, a reduction of 5.66% is expected (Fig. 6k), mainly concentrated in the middle reaches and middle and lower reaches of the Yellow River in China (Fig. 6j, refer to red color). Unlike other species, *V. romanetii* has an Expansion-Contraction Ratio of zero for its suitable areas, indicating it is at a critical threshold, further research and adaptive farming strategies may be necessary to ensure the viability of the crop. In general, under the impending climate change, eight of the ten species exhibit significant adaptive capabilities, these species stand out for future agricultural efforts due to their positive growth trajectories and resilience to anticipated climate changes. Their favorable rate of expansion contrasts with *V. amurensis*, which is facing contraction. This emphasizes their inherent adaptability and potential to flourish in the face of a changing climate.

**Fig. 6.**
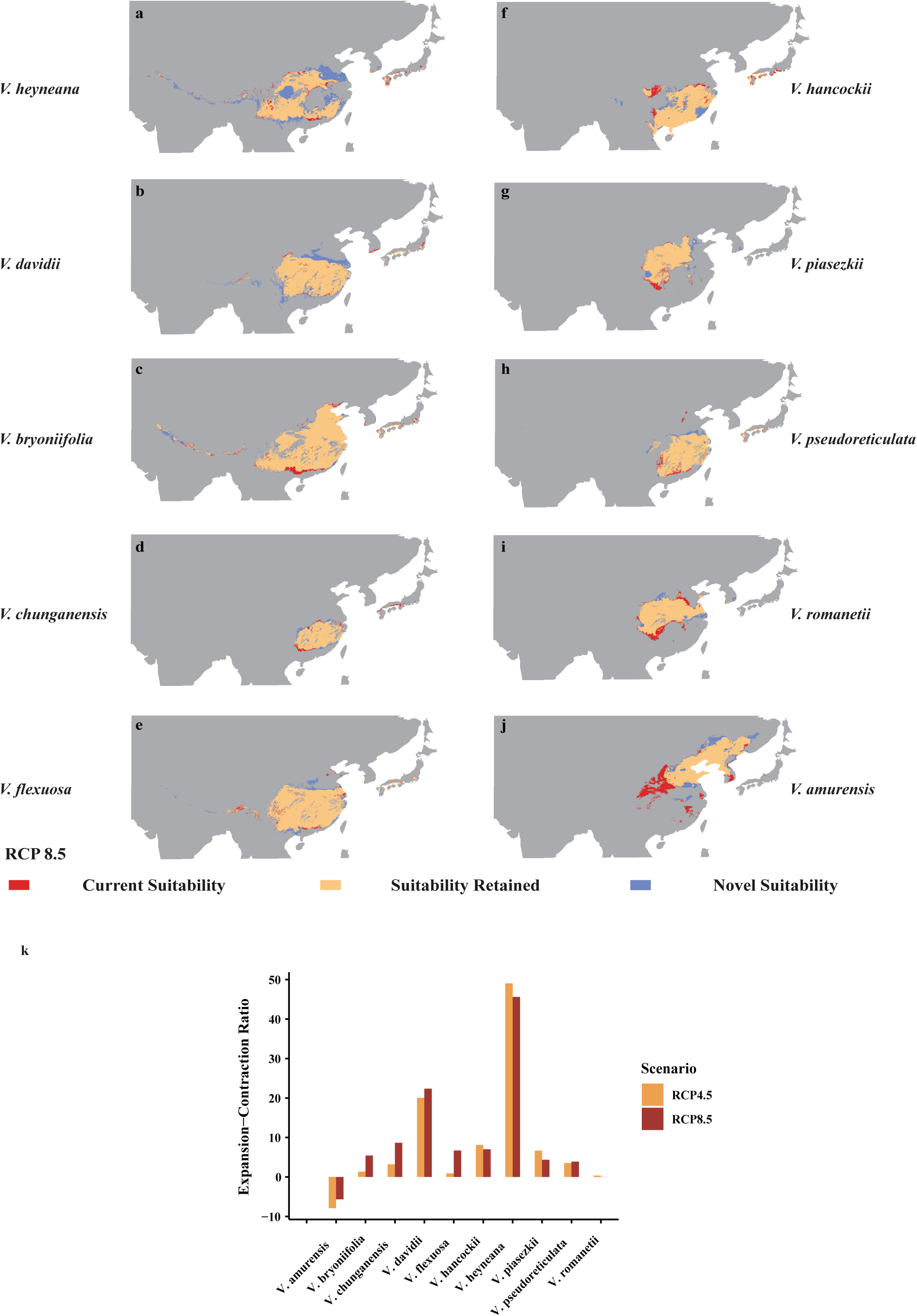
Maxent model projected bioclimatic suitability for East Asian wild grapes (a) *V. heyneana*, (b) *V. davidi*, (c) *V. bryoniifolia*, (d) *V. chunganensis*, (e) *V. flexuosa*, (f) *V. hancockii*, (g) *V. piasezkii*, (h) *V. pseudoreticulata*, (i) *V. romanetii*, (j) *V. amurensis*, with future projections for the years 2081–2100 under the RCP 8.5 climatic scenarios. (k) Net suitability change for the distribution of East Asian wild grapes. Bar plots show Expansion-Contraction Ratio of change in area suitable for grape-growing regions projected by maxent model for RCP 8.5 (red) and RCP 4.5 (yellow).

## Discussion

### Grapevine and its wild relatives in response to climate change (comparing grape and its wild relatives)

A decline in crop suitability caused by global climate change could lead to significant economic and conservation impacts, with variable responses depending on crop type, region, and adaptation strategies, highlighting the urgent need for targeted agricultural practices to ensure food security (Zhou et al., 2014; Rezaei et al., 2023; Hosokawa et al., 2023; Mohammadi et al., 2023; Farooq et al., 2023; Magon et al., 2023). Furthermore, climate change, characterized by rising temperatures and changing precipitation patterns, presents a significant threat to the global wine industry (Hannah et al., 2013; Cabré et al., 2020; Zhang et al., 2024; Van et al., 2024). However, a comprehensive evaluation of the impacts of future climate change on cultivated grape and its wild relative remains necessary. Our results further underscore the profound implications of climate change on viticulture and suggest that the potential expansion of wild grape growth in North America as climate conditions evolve presents both opportunities and challenges (Fig. 5). In contrast, under the RCP 8.5 scenario, which projects a significant increase in greenhouse gas emissions, we predict a substantial reduction in the areas suitable for wine grape cultivation. This change could make some traditional grape cultivation regions unsuitable for existing grape varieties, potentially leading to losses of 15.04×10^5^ km² for wine grapes and 13.26×10^5^ km² for table grapes (Table S1). This change may render traditional grape-growing regions unsuitable for current varieties, requiring the development of new varieties that can adapt to climate change. Furthermore, we identify an expansion of the suitable areas for table grapes beyond that for wine grapes, suggesting that table grapes may be more resilient to the severe weather conditions expected in the future (Fig. 3, Table S1). This unexpected identify may indicate that table grapes have a greater potential to thrive under expected environmental stresses compared to wine grapes in the face of climate change. Compared to cultivated grapes, wild grapes may have a greater ability to adapt to future climate change. North American and East Asian wild grapes, in particular, may exhibit even greater adaptability than European ones, such as *V. sylvestris* (Fig. 4). For instance, wild grapes in North America and East Asia have demonstrated remarkable adaptability to extreme temperature changes (Fig. 5k and 6k). Their ability to survive and thrive in warmer climates can be attributed to the genetic diversity shaped by long-term natural selection (Fig. 5 and 6). The genetic diversity and stress tolerance traits in these wild grapes could be invaluable resources for breeding programs aimed at developing new cultivars better able to withstand the environmental pressures of climate change. As the global climate continues to change, this insight could significantly impact the grape industry, potentially leading to a repositioning of table and wild grapes to ensure economic viability and the ability of agriculture against future climate uncertainties. The urgency to adapt agricultural practices and the need for innovation in grape breeding, highlights the critical need to ensure the sustainability and evolution of the viticulture industry in the face of global warming.

In response to climate challenges, strategic crop switching and relocation in the United States may reduce the decline in agricultural profits by up to 50% under the RCP 8.5 scenario (Rising et al., 2020). Crop migration is a crucial climate adaptation strategy that can protect yields of major crops, although its success depends on overcoming socio-economic and environmental obstacles (Sloat et al., 2020). Globally, optimizing cropland distribution through modeling can reduce the environmental impact of agriculture whil<u>e</u> maintaining food production. Innovative adaptation measures, such as Ethiopian farmers transitioning to drought-resistant perennial crops like enset, are enhancing food security in drought-prone regions (Beyer et al., 2023; Chase et al., 2023). Actually, these strategies emphasize the importance of proactive and innovative measures in agriculture to address the complex climate change challenges and ensure the sustainability of both agricultural profits and environmental health. However, these studies also highlight the limitations and challenges of these strategies, including the need for further adaptation measures, uncertainty about long-term sustainability, and potential environmental costs associated with such large-scale shifts in agricultural practices. In summary, while crop switching and migration present viable short- and medium-term solutions to climate change pressures on agriculture, they also emphasize the need for a more comprehensive and sustainable approach to agricultural adaptation that addresses economic, environmental, and regional complexities.

### Wild *Vitis* genetic resources for breeding grapes and its rootstocks with climate resilience

In the face of climate change, researchers have identified a valuable genetic resource of CWRs and plants used as rootstocks to enhance the resilience and adaptability of domesticated crops (Rogiers et al., 2022). These wild plants are closely related to cultivated crops and are increasingly recognized as valuable genetic resources that enhance the ability of crops to cope with various stresses and agricultural sustainability challenges. Recent advances in genomics and high-throughput techniques are accelerating the development of climate-resilient crops with enhanced stress tolerance and nutritional value (Satori et al., 2022). This includes harnessing the genetic potential of CWRs to create new varieties and polyploids, which is essential for adapting to future climates (Pourkheirandish et al., 2020; Hohenlohe et al., 2021). In summary, integrating CWRs into crop improvement programs, supported by genomics and conservation strategies, is crucial for building resilience against climate change. This approach not only enhances our crops’ genetics but also ensures the long-term sustainability of global food systems, safeguarding nutritious food for future generations (Bohra et al., 2022; Coyne et al., 2020). Our research provides a comprehensive analysis of the adaptability of various wild grape species, including European *V. sylvestris*, and identifies that specific species of East Asian and North American wild grapes exhibit exceptional abilities to adapt to changing climate patterns, indicating their potential as a genetic bridge toward a more resilient and adaptable form of grape range. Wild grapes in North America, particularly *V. rotundifolia* and *V. labrusca*, are predicted to undergo substantial increases in suitable distribution areas under different climate change scenarios, with *V. rotundifolia* leading the expansion, with projections of 227% under RCP 4.5 scenario, and 205% under RCP 8.5 scenario (Fig. 5k). In addition to the well-known *V. labrusca* and *V. rotundifolia*, other species such as *V. arizonica*, *V. monticola*, and *V. vulpina* have also demonstrated exceptional adaptability. Similarly, East Asian wild grapes such as *V. heyneana* are predicted to expand their suitable range, while *V. davidii* shows a unique response to climate scenarios, to be more specific, the expansion-contraction ratio of the change in area suitable for *V. davidii* regions, as projected by the RCP 8.5, is higher than that projected by the RCP 4.5 Fig. 6k). Furthermore, other species such as *V. bryoniifolia*, *V.* (*chunganensis*, *V. flexuosa*, *V. hancockii*, *V. piasezkii*, and *V. pseudoreticulata* have demonstrated strong adaptability, making them promising candidates for future distribution. Previous researchers had found that studying these species, including six wild grapes in the southwestern United States and comparing them with Chinese wild grapes (*V. davidii*), and revealed that these species have evolved genetic networks and disease resistance genes that enhance their ability to adapt to environmental stresses such as high temperatures or drought (Zhang et al., 2019; Morales-Cruz et al., 2021). Overall, these wild species demonstrate genetic resilience and potential resistance to environmental stresses like drought and heat, qualities essential for maintaining the sustainability of grape horticulture practices. Moreover, these species have potential in developing rootstocks that can enhance the adaptability of cultivated grapes and help produce high-quality grapes under climate change. The results provide a new insight into grape-environment interactions, ultimately supporting the selection of suitable rootstocks.

## Conclusion

Our study employed meteorological factors and the maxent model to predict shifts in suitable agricultural regions for wild grapes and cultivated grapes under future climatic conditions. The research underscores the enhanced adaptability of North American wild grapes and East Asian wild grapes in the face of projected climate changes. Using the maxent model, we identified that these specific wild grape species are likely to expand their suitable habitats in future climates, which provides them with sensitivity and adaptability to changing environmental conditions, allowing them to survive and thrive in warmer climates. This insight is significant for understanding the geographical distribution and ecological resilience of grape species. Furthermore, our research establishes a foundation for further exploration into resistance gene mining and rootstock improvement. This is essential for addressing climate change challenges and ensuring the sustainable development of the grape industry. In summary, our study provides a scientific basis for the planning of grape cultivation regions and offers a novel perspective on the conservation and utilization of grape genetic resources. Future research will focus on genomic analysis of these wild grapes and the application of their resistance genes in the improvement of cultivated varieties.

## Materials and Methods

### Data Collection and Analysis of *V.* L. Distribution

The distribution data of *V.* L. were collected from GBIF (https://www.gbif.org/), VIVC (https://www.vivc.de/), and related literature (Puga et al., 2022). Flora of China, and Flora of North America was used to check the distribution locations with clear records of longitude and latitude. It will supplement some distribution data with missing geographical coordinate information and eliminate locations that cannot be accurately located due to missing geographic coordinate information or unclear descriptions. Distribution data that clearly deviated from geographic coordinate information and administrative place name records were also eliminated. Finally, 5,662 effective distribution sites were obtained (2,109 for cultivated grapes and 3,553 for wild grapes). The obtained data, which retains only the longitude, latitude, and species name, is saved in CSV format and used as the source for establishing the species distribution model (SDM).

### Source and processing of climate data

Current and future climate data were derived from the WorldClim database (https://www.worldclim.org/), with a spatial resolution of 2.5 arc-minute grid (geospatial resolution of approximately 5 km), and includes 19 bioclimatic variables (Bio1∼Bio19). Future climate data (2081-2100) uses the EC-Earth3-Veg atmospheric circulation model to evaluate the potential impact of various policies and development paths on climate change by simulating different Shared Socioeconomic Pathways (SSP) emission scenarios.

### Model building, optimization and evaluation

Apply the processed distribution data and environmental factors to the maxent model for analysis, models were fit by applying the machine-learning algorithm maxent 3.4.3 (Phillips et al., 2006). The parameter settings of the initial model are: 25% distribution points are used as the random test percentage in the model, the default maximum number of background points is 10,000, running 10 model replicates, and Jackknife is used to calculate the contribution rate of environmental factors to explore the relationship between the suitable zone and relationship with environment variables. Model construction accuracy is determined by the area under the curve (AUC), with values ranging from 0 to 1. Higher AUC values indicate better model fit, higher construction accuracy, and greater credibility. An AUC value of 0.5-0.6 indicates failure in model construction, 0.6-0.7 indicates poor simulation effect, 0.7-0.8 indicates average simulation effect, 0.8-0.9 indicates good simulation effect, and 0.9-1 indicates excellent simulation effect (Konowalik., 2021).

### ArcGIS and R for spatial data analysis and visualization

We utilize the SDMtoolbox 2.0 extension in ArcGIS 10.8.2 to analyze areas of expansion and contraction, effectively predicting and addressing the potential impacts of climate change on the geographical distribution of grape populations (Zhang et al., 2017; Hazarika et al., 2023). In addition, we are conducting an in-depth analysis of the ratios of expansion and contraction areas for grapes under different climate scenarios, aiming to develop more accurate adaptation strategies, and are using RStudio-2024.04.2+764 to create bar charts illustrating the percentages.

## Acknowledgements

This work was supported by the National Natural Science Foundation of China (No. 32300191; 32372662). We are also particularly grateful for the services of the High-Performance Computing Cluster and experimental sites in the Agricultural Genomics Institute at Shenzhen, Chinese Academy of Agricultural Sciences.

## Data availability

All data needed to evaluate the conclusions in this paper are presented in the paper and the supplementary information.

## Reference

1. Adhikari, U., Nejadhashemi, A. P., & Woznicki, S. A. Climate change and eastern Africa: a review of impact on major crops. Food and Energy Security 4, 110–132 10.1002/fes3.61 (2015).

2. Aguirre□Liguori, J. A., Morales□Cruz, A., & Gaut, B. S. Evaluating the persistence and utility of five wild *Vitis* species in the context of climate change. Molecular Ecology 31, 6457–6472 10.1111/mec.16715 (2022).

3. Beyer, R.M., Hua, F., Martin, P.A., et al. Relocating croplands could drastically reduce the environmental impacts of global food production. Communications Earth & Environment 3, 49 10.1038/s43247-022-00360-6 (2022).

4. Bohra, A., Kilian, B., Sivasankar, S., et al. Reap the crop wild relatives for breeding future crops. Trends in Biotechnology 40, 412–431 10.1016/j.tibtech.2021.08.009 (2022).

5. Bigolin, T.; Talamini, E. Impacts of Climate Change Scenarios on the Corn and Soybean Double-Cropping System in Brazil. Climate 12, 42 10.3390/cli12030042 (2024).

6. Brett R. Scheffers et al. The broad footprint of climate change from genes to biomes to people. Science 354, aaf 7671 https://www.science.org/doi/abs/10.1126/science.aaf7671 (2016).

7. Cabré, F., Nuñez, M. Impacts of climate change on viticulture in Argentina. Regional Environmental Change 20, 12 10.1007/s10113-020-01607-8 (2020).

8. Caffarra, Amelia, et al. Modelling the impact of climate change on the interaction between grapevine and its pests and pathogens: European grapevine moth and powdery mildew. Agriculture, Ecosystems & Environment 148, 89–101 10.1016/j.agee.2011.11.017 (2012).

9. Chase, R. R., Büchi, L., Rodenburg, J. et al. Smallholder farmers expand production area of the perennial crop enset as a climate coping strategy in a drought□prone indigenous agrisystem. Plants People Planet 5, 254–266 10.1002/ppp3.10339 (2023).

10. Cochetel, N., Minio, A., Guarracino, A., et al. A super-pangenome of the North American wild grape species. Genome Biology 24, 290 10.1186/s13059-023-03133-2 (2023).

11. Cornelis van Leeuwen, Agnes Destrac-Irvine. Modified grape composition under climate change conditions requires adaptations in the vineyard. OENO One 51, 147–154 https://oeno-one.eu/article/view/1647 (2017).

12. Coyne, C. J., Kumar, S., Wettberg, E. J. B. V. et al. Potential and limits of exploitation of crop wild relatives for pea, lentil, and chickpea improvement. Legume Science 2, e36 10.1002/leg3.36 (2020).

13. David B. Lobell., et al. Climate trends and global crop production since 1980. Science 333, 616–620 https://www.science.org/doi/10.1126/science.1204531 (2011).

14. De Orduna, Ramon Mira. Climate change associated effects on grape and wine quality and production. Food Research International 43, 1844–1855 10.1016/j.foodres.2010.05.001 (2010).

15. Dhaliwal, D.S., Williams, M.M. Evidence of sweet corn yield losses from rising temperatures. Scientific Reports 12, 18218 10.1038/s41598-022-23237-2 (2022).

16. Farooq, A., Farooq, N., Akbar, H. et al. A critical review of climate change impact at a global scale on cereal crop production. Agronomy 13, 162 10.3390/agronomy13010162 (2023).

17. Fernie, A. R., & Yan, J. De novo domestication: an alternative route toward new crops for the future. Molecular plant 12, 615–631 10.1016/j.molp.2019.03.016 (2019).

18. Hannah L, Roehrdanz PR, Ikegami M. et al. Climate change, wine, and conservation. Proceedings of the National Academy of Sciences of the United States of America 110, 6907–6912 10.1073/pnas.1210127110 (2013).

19. Hazarika A, Deka J R, Nath P C, et al. Modelling habitat suitability of the critically endangered Agarwood (*Aquilaria malaccensis*) in the Indian East Himalayan region. Biodiversity and Conservation 32, 4787–4803 10.1007/s10531-023-02727-3 (2023).

20. Hohenlohe, P. A., Funk, W. C., & Rajora, O. P. Population genomics for wildlife conservation and management. Molecular Ecology 30, 62–82 10.1111/mec.15720 (2021).

21. Hosokawa, N., Doi, Y., Kim, W., et al. Contrasting area and yield responses to extreme climate contributes to climate-resilient rice production in Asia. Scientific Reports 13, 6219 10.1038/s41598-023-33413-7 (2023).

22. Konowalik, K., Nosol, A. Evaluation metrics and validation of presence-only species distribution models based on distributional maps with varying coverage. Scientific Reports 11, 1482 10.1038/s41598-020-80062-1 (2021).

23. Li, T., Yang, X., Yu, Y., et al. Domestication of wild tomato is accelerated by genome editing. Nature Biotechnology 36, 1160–1163 10.1038/nbt.4273 (2018).

24. Lin, Y., Chen, Y., Wang, H. et al. Global potential distributions and conservation status of rice wild relatives. Plants, People, Planet. 10.1002/ppp3.10522 (2024).

25. Ma, Z. Y., Nie, Z. L., Liu, X. Q. et al. Phylogenetic relationships, hybridization events, and drivers of diversification of East Asian wild grapes as revealed by phylogenomic analyses. Journal of Systematics and Evolution 61, 273–283 10.1111/jse.12918 (2023).

26. Magon, G., De Rosa, V., Martina, M., et al. Boosting grapevine breeding for climate-smart viticulture: from genetic resources to predictive genomics. Frontiers in Plant Science, 14 1293186. 10.3389/fpls.2023.1293186 (2023).

27. Mohammadi, S., Rydgren, K., Bakkestuen, V. et al. Impacts of recent climate change on crop yield can depend on local conditions in climatically diverse regions of Norway. Scientific Reports 13, 3633 10.1038/s41598-023-30813-7 (2023).

28. Morales-Castilla I, García de Cortázar-Atauri I, Cook BI. et al. Diversity buffers winegrowing regions from climate change losses. Proceedings of the National Academy of Sciences of the United States of America 117, 2864–2869 10.1073/pnas.1906731117 (2020).

29. Morales-Cruz, A., Aguirre-Liguori, J.A., Zhou, Y., et al. Introgression among North American wild grapes (*Vitis*) fuels biotic and abiotic adaptation. Genome Biology 22, 254 10.1186/s13059-021-02467-z (2021).

30. Morales-Cruz, A., Aguirre-Liguori, J., Massonnet, M. et al. Multigenic resistance to Xylella fastidiosa in wild grapes (Vitis sps.) and its implications within a changing climate. Communications Biology 6, 580 10.1038/s42003-023-04938-4 (2023)

31. Muller, C., Cramer, W., Hare, W. L. et al. Climate change risks for African agriculture. Proceedings of the National Academy of Sciences of the United States of America 108, 4313–4315 10.1073/pnas.1015078108 (2011).

32. Tim Wheeler, Joachim von Braun. Climate change impacts on global food security. Science 341, 508–513 https://www.science.org/doi/abs/10.1126/science.1239402 (2013).

33. Petitpierre, B., Arnold, C., Phelps, L. N. et al. A tale of three vines: current and future threats to wild Eurasian grapevine by vineyards and invasive rootstocks. Diversity and Distributions 29, 1594–1608 10.1111/ddi.13780 (2023).

34. Phillips, S. J., Anderson, R. P., & Schapire, R. E. Maximum entropy modeling of species geographic distributions. Ecological Modelling 190, 231–259 10.1016/j.ecolmodel.2005.03.026 (2006).

35. Pironon, S., Etherington, T. R., Borrell, J. S. et al. Potential adaptive strategies for 29 sub-Saharan crops under future climate change. Nature Climate Change 9, 758–763 10.1038/s41558-019-0585-7 (2019).

36. Porch, T. G., Beaver, J. S., Debouck, D. G. et al. Use of wild relatives and closely related species to adapt common bean to climate change. Agronomy 3, 433–461 10.3390/agronomy3020433 (2013).

37. Pourkheirandish, M., Golicz, A. A., Bhalla, P. L. et al. Global role of crop genomics in the face of climate change. Frontiers in Plant Science 11, 922 10.3389/fpls.2020.00922 (2020).

38. Rahimi, O., Ohana Levi, N., Brauner, H. et al. Demographic and ecogeographic factors limit wild grapevine spread at the southern edge of its distribution range. Ecology and Evolution 11, 6657–6671 10.1002/ece3.7519 (2021).

39. Renzi, J. P., Coyne, C. J., Berger, J. et al. How could the use of crop wild relatives in breeding increase the adaptation of crops to marginal environments?. Frontiers in Plant Science 13, 886162 10.3389/fpls.2022.886162 (2022).

40. Rezaei, E.E., Webber, H., Asseng, S., et al. Climate change impacts on crop yields. Nature Reviews Earth & Environment 4, 831–846 10.1038/s43017-023-00491-0 (2023).

41. Rising, J., Devineni, N. Crop switching reduces agricultural losses from climate change in the United States by half under RCP 8.5. Nature Communications 11, 4991 10.1038/s41467-020-18725-w (2020).

42. Rogiers, S. Y., Greer, D. H., Liu, Y. et al. Impact of climate change on grape berry ripening: An assessment of adaptation strategies for the Australian vineyard. Frontiers in Plant Science 13, 1094633 10.3389/fpls.2022.1094633 (2022).

43. Satori, D., Tovar, C., Faruk, A. et al. Prioritising crop wild relatives to enhance agricultural resilience in sub□Saharan Africa under climate change. Plants, People, Planet 4, 269–282 10.1002/ppp3.10247 (2022).

44. Sloat, L.L., Davis, S.J., Gerber, J.S., et al. Climate adaptation by crop migration. Nature Communications 11, 1243 10.1038/s41467-020-15076-4 (2020).

45. Vadez, V., Berger, J. D., Warkentin, T. et al. Adaptation of grain legumes to climate change: a review. Agronomy for Sustainable Development 32, 31–44 10.1007/s13593-011-0020-6 (2012).

46. Van Leeuwen, C., Destrac-Irvine, A., Dubernet, M. et al. An update on the impact of climate change in viticulture and potential adaptations. Agronomy 9, 514 10.3390/agronomy9090514 (2019).

47. Van Leeuwen, C., Sgubin, G., Bois, B., et al. Climate change impacts and adaptations of wine production. Nature Reviews Earth & Environment 5, 258–275 10.1038/s43017-024-00521-5 (2024).

48. Waha K, Van Wijk M T, Fritz S., et al. Agricultural diversification as an important strategy for achieving food security in Africa. Global Change Biology 24, 3390–3400 10.1111/gcb.14158 (2018).

49. Wang N, Cao S, Liu Z., et al. Genomic conservation of crop wild relatives: A case study of citrus. PLoS Genetics 19, e1010811. 10.1371/journal.pgen.1010811 (2023).

50. Wang Z, Wong D C J, Wang Y., et al. GRAS-domain transcription factor PAT1 regulates jasmonic acid biosynthesis in grape cold stress response. Plant Physiology 186, 1660–1678 10.1093/plphys/kiab142 (2021).

51. Xiao, H., Liu, Z., Wang, N. et al. Adaptive and maladaptive introgression in grapevine domestication. Proceedings of the National Academy of Sciences of the United States of America 120, e2222041120 10.1073/pnas.2222041120 (2023).

52. Yadav, S. S. Crop wild relatives and climate change (p. 400). R. Redden, N. Maxted, M. E. Dulloo, L. Guarino, & P. Smith (Eds.). Hoboken, NJ, USA: Wiley-Blackwell. https://onlinelibrary.wiley.com/doi/book/10.1002/9781118854396 (2015).

53. Yang Dong et al. Dual domestications and origin of traits in grapevine evolution. Science 379, 892–901 https://www.science.org/doi/full/10.1126/science.add8655 (2023).

54. Yi Yang et al. Climate change exacerbates the environmental impacts of agriculture. Science 385, eadn3747 https://www.science.org/doi/abs/10.1126/science.adn3747 (2024).

55. You, Y., Yu, J., Nie, Z., et al. Transition of survival strategies under global climate shifts in the grape family. Nat. Plants 10, 1100–1111 10.1038/s41477-024-01726-8 (2024).

56. Zhang F, Long R, Ma Z. et al. Evolutionary genomics of climatic adaptation and resilience to climate change in alfalfa. Molecular Plant, 10.1016/j.molp.2024.04.013 (2024).

57. Zhang, H., Mittal, N., Leamy, L. J. et al. Back into the wild—Apply untapped genetic diversity of wild relatives for crop improvement. Evolutionary applications 10, 5–24 10.1111/eva.12434 (2017).

58. Zhang L, Liu S, Sun P., et al. Consensus forecasting of species distributions: the effects of niche model performance and niche properties. Public Library of Science 10, e0120056 10.1371/journal.pone.0120056 (2015).

59. Zhang T, Peng W, Xiao H., et al. Population genomics highlights structural variations in local adaptation to saline coastal environments in woolly grape. Journal of Integrative Plant Biology. 10.1111/jipb.13653 (2024).

60. Zhang, Y., Yao, JL., Feng, H., et al. Identification of the defense-related gene VdWRKY53 from the wild grapevine *Vitis davidii* using RNA sequencing and ectopic expression analysis in Arabidopsis. Hereditas 156, 14 10.1186/s41065-019-0089-5 (2019).

61. Zhou Y, Zhang L, Liu J., et al. Climatic adaptation and ecological divergence between two closely related pine species in Southeast China. Molecular Ecology 23, 3504–3522 10.1111/mec.12830 (2014).

62. Zhou Y, Massonnet M, Sanjak J S., et al. Evolutionary genomics of grape (*Vitis vinifera* ssp. *vinifera*) domestication. Proceedings of the National Academy of Sciences of the United States of America 114, 11715–11720 10.1073/pnas.1709257114 (2017).

