## Supplementary material for "The dynamics of wild *Vitis* species in response to climate change facilitate the breeding of grapevine and its rootstocks with climate resilience": fig s1-s5

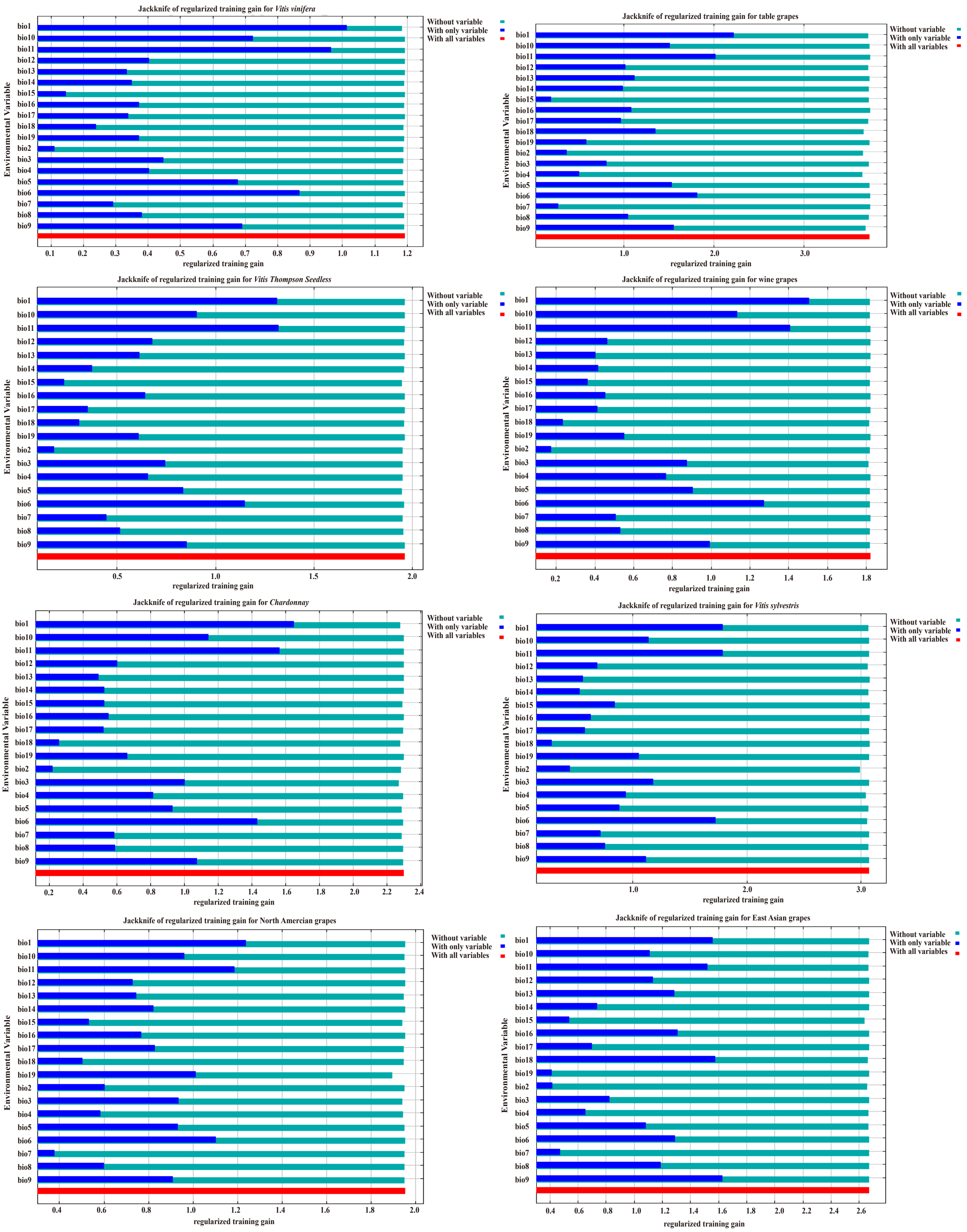


**Fig. S1. Evaluation of environmental factors byJakknife method**

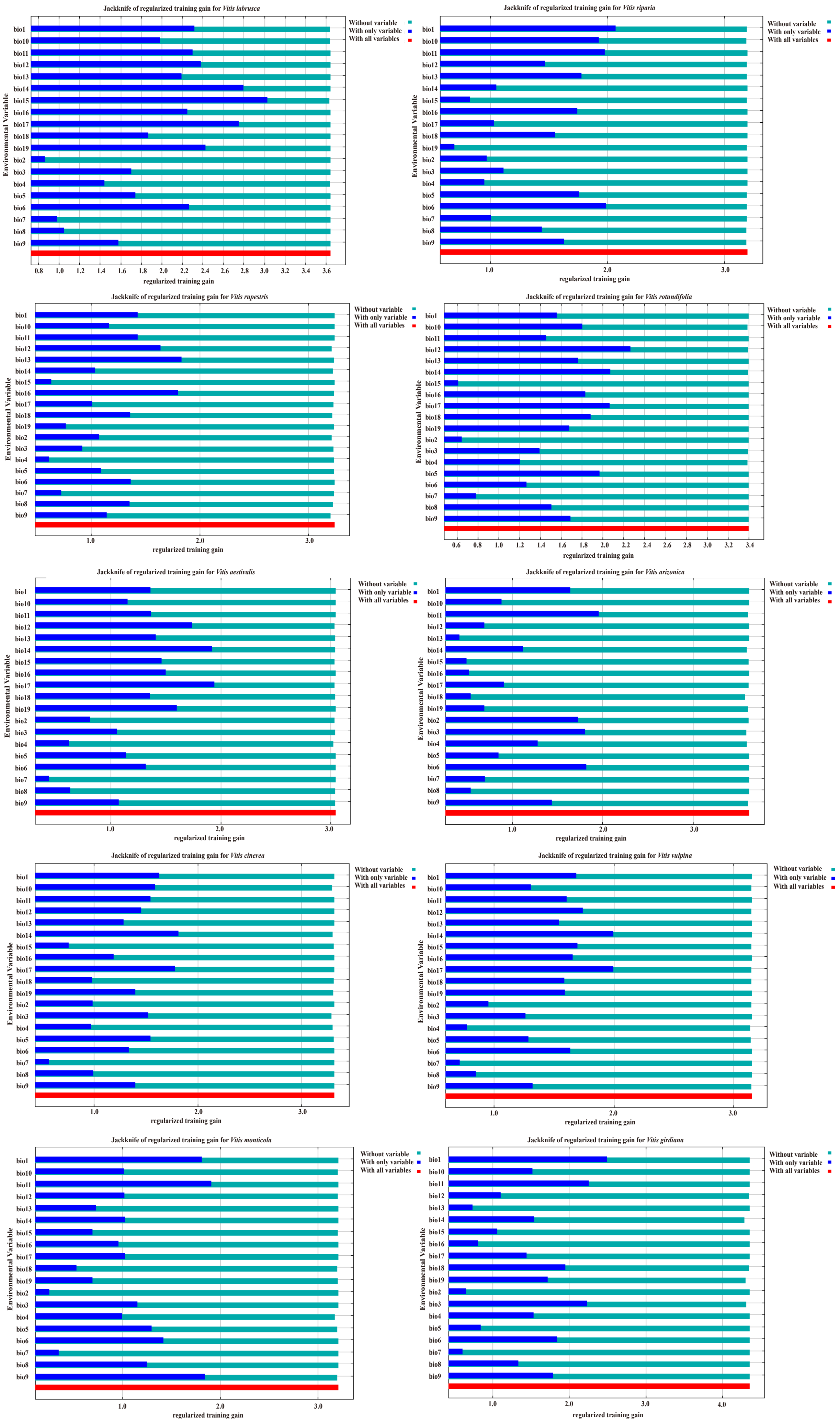


**Fig. S1. Evaluation of environmental factors byJakknife method**

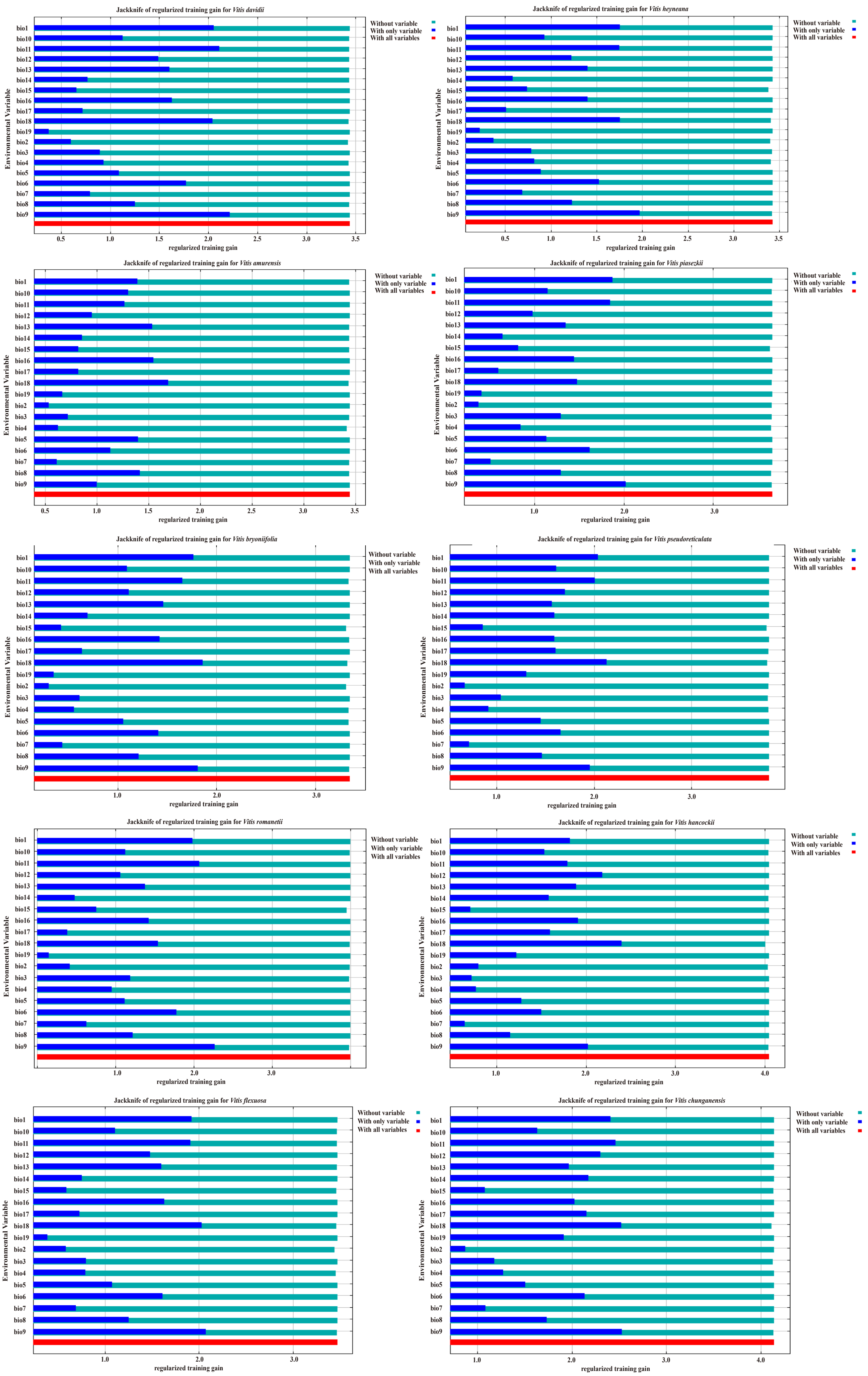


**Fig. S1. Evaluation of environmental factors byJakknife method**

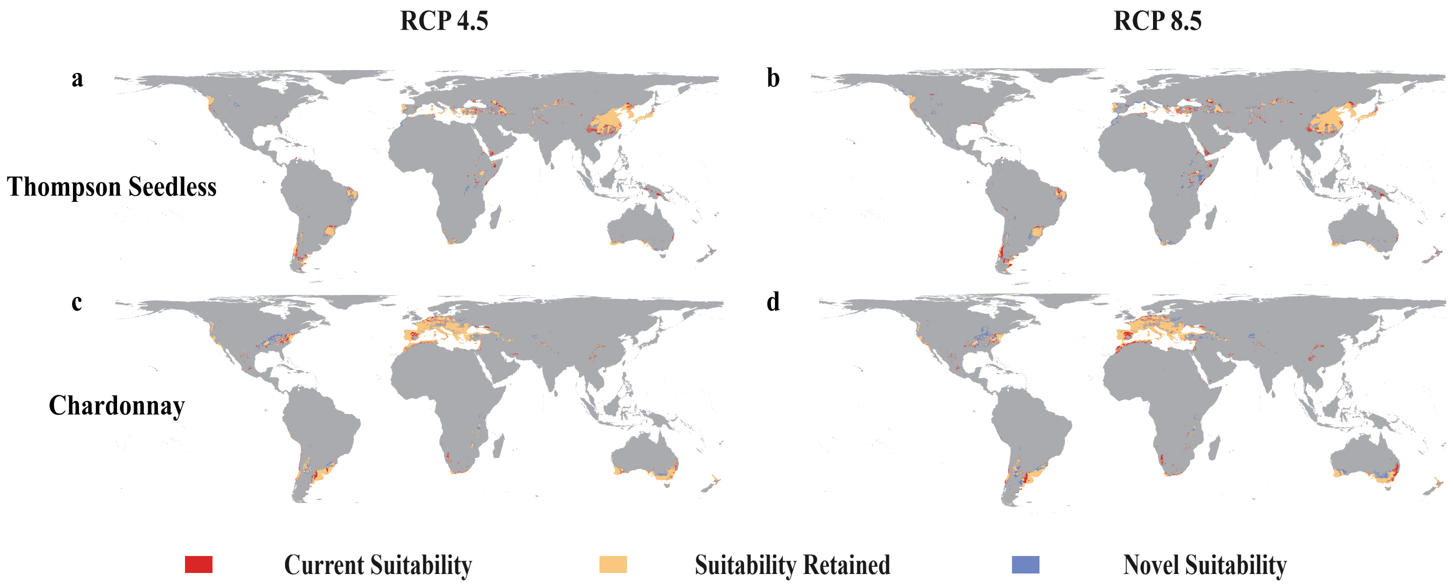


**Fig. S2. Maxent model projected bioclimatic suitability for Thompson Seedless and Chardonnay (a-b) Thompson Seedless, (c-d) Chardonnay. Future bioclimatic suitability projections (2081–2100), under the RCP 4.5 and RCP 8.5 climatic scenario.**

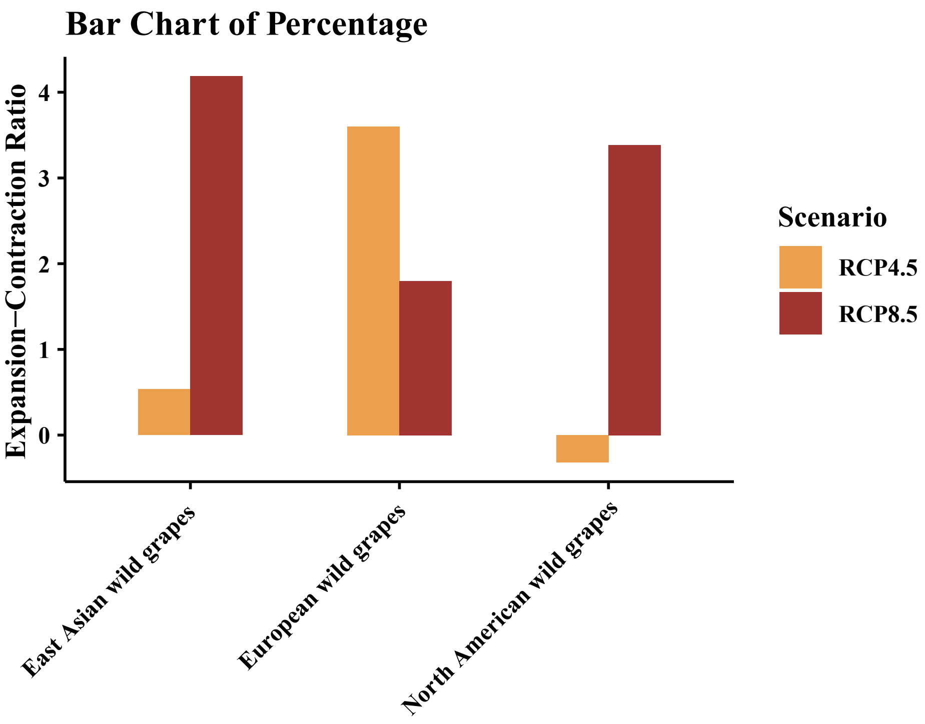


**Fig. S3. Net suitability change for the distribution of European wild grapes (Vitis sylvestris), North American wild grapes, and East Asian wild grapes. Bar plots show Expansion-Contraction Ratio of change in area suitable for grape-range projected by maxent model for RCP 8.5 (red) and RCP 4.5 (yellow).**

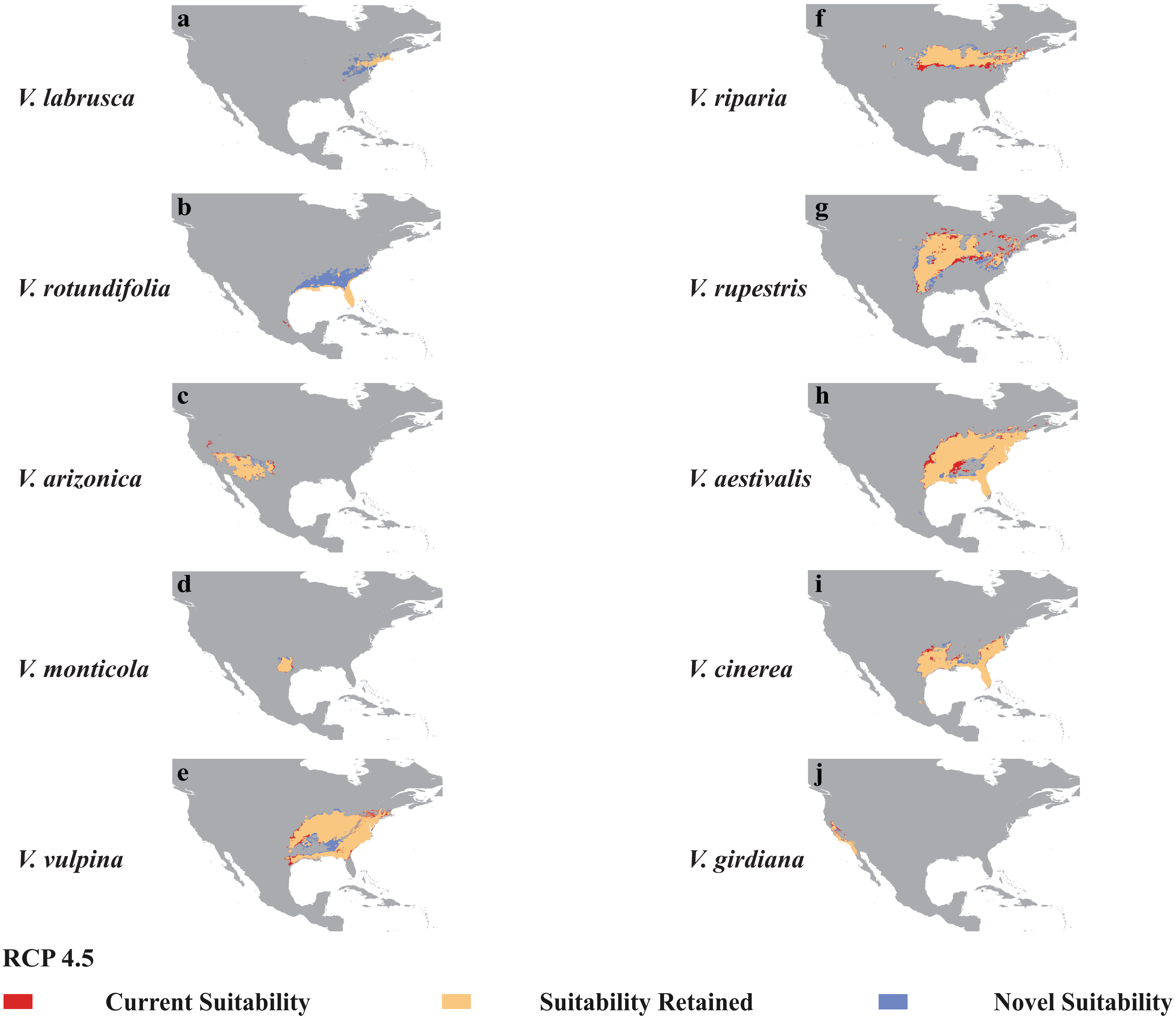


**Fig. S4. Maxent model projected bioclimatic suitability for North American wild grapes (a) *V. labrusca*, (b) *V. rotundifolia*, (c) *V. arizonica*, (d) *V. monticola*, (e) *V. vulpina*, (f) *V. riparia*, (g) *V. rupestris*, (h) *V. aestivalis*, (i) *V. cinerea*, (j) *V. girdiana*. Future bioclimatic suitability projections (2081–2100), under the RCP 4.5 climatic scenario.**

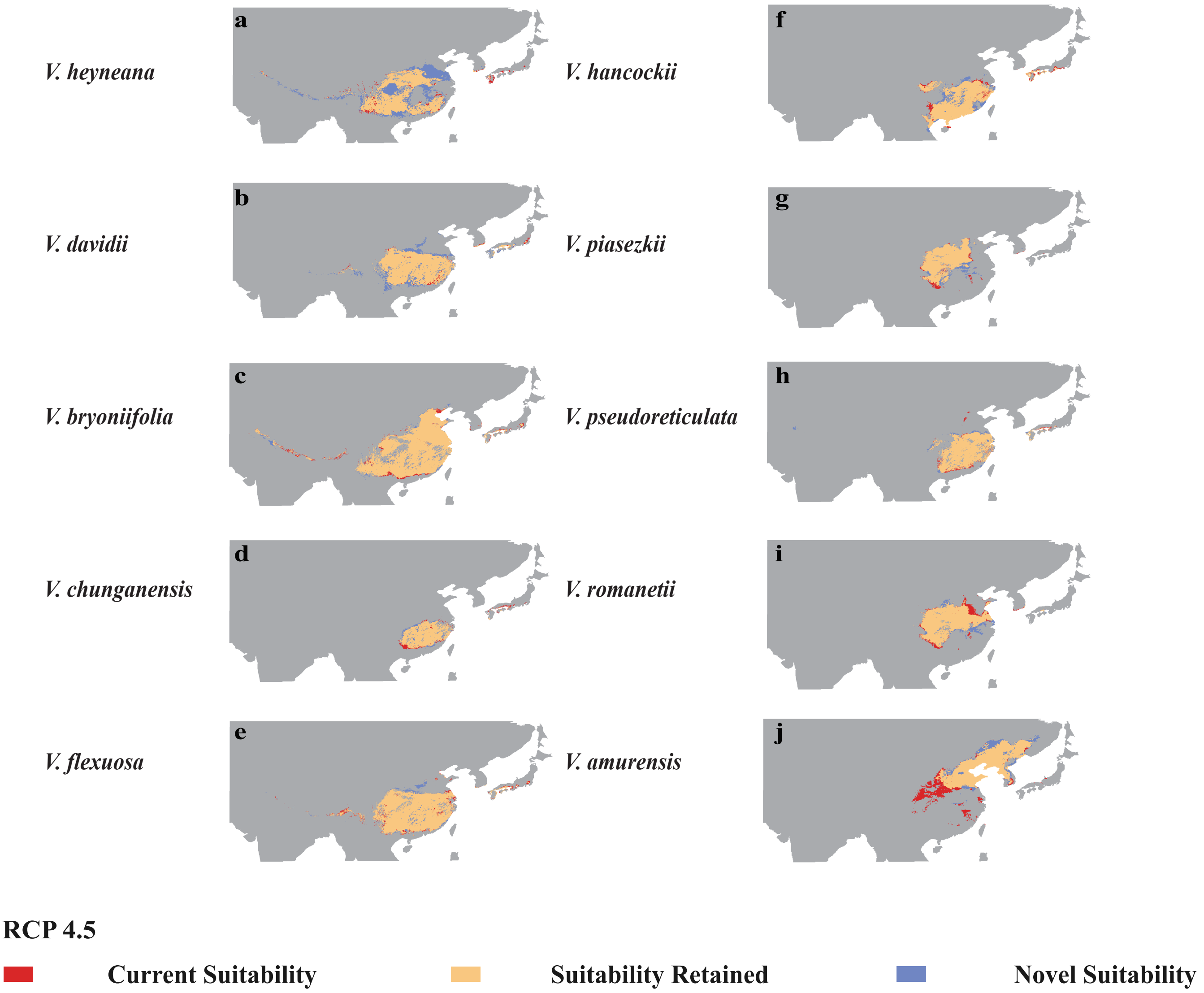


**Fig. S5. Maxent model projected bioclimatic suitability for North American wild grapes (a) *V. heyneana*, (b) *V. davidii*, (c) *V. bryoniifolia*, (d) *V. chunganensis*, (e) *V. flexuosa*, (f) *V. hancockii*, (g) *V. piasezkii*, (h) *V. pseudoreticulata*, (i) *V.* romanetii, (j) *V.* amurensis. Future bioclimatic suitability projections (2081–2100), under the RCP 4.5 climatic scenario.**
